# Visuomotor activity associated with conditioned hand movements in premotor cortex of blindsight macaques

**DOI:** 10.1101/2023.02.17.529035

**Authors:** Yusuke Yamamoto, Saya Kitazume, Jun Takahashi, Hirotaka Onoe, Reona Yamaguchi, Tadashi Isa

## Abstract

Some patients with damage to the primary visual cortex (V1) can respond to visual stimuli in their affected visual field, even though they report loss of visual awareness. This phenomenon is termed blindsight. To clarify the visuomotor transformation for hand movement control in the blindsight condition, we conducted multichannel electrocorticographical recordings from the ipsilesional frontal cortex including the frontal eye field, premotor, primary motor, and somatosensory areas during a delayed conditioned two-alternative forced choice manual response task with arbitrary assignment of push-pull manual responses in relation to the visual target cue locations before and after unilateral V1 lesioning in macaque monkeys. Before lesioning, the activity in the dorsal premotor cortex (PMd) showed short latency task-related θ ~ α band responses that were significantly higher in the successful trials than in the error/miss trials at approximately 200 ms after target cue onset. At 2–3 months after lesioning, the monkeys regained a >80% success rate in response to appearance of the target cue in the lesion-affected visual field. At this stage, similar task-related θ ~ α band target cue responses were observed in the PMd. Further, Granger causality from the PMd to the primary motor cortex was enhanced in the θ ~ α, γ, and high-γ bands during the delay period of the task. These results suggested that the PMd plays a crucial role in mediating the visual signals for execution of hand movements in blindsight monkeys.

**Significant Statement:** Blindsight is a curious phenomenon in which the patients with damage to the primary visual cortex can still respond to visual stimulus in their blinded visual field despite loss of awareness. Previously we clarified the critical circuits for visually guided saccadic eye movements in blindsight macaques, however, those for the control of hand movements are still unclear. Here, we have recorded the multichannel electrocorticography in the frontal cortices during the conditioned manual response task and found that the dorsal premotor cortex exhibits task-related θ ~ α band target cue responses at approximately 200 ms after the visual cue onset, similarly to the intact state. This is the first report of electrophysiological recordings in the frontal lobe of the blindsight subjects.

## Introduction

Damage to the primary visual cortex (V1) impairs subjective visual awareness; however, some patients exhibit behavioral responses to visual stimuli in their affected visual field when they are forced to perform by guessing (Pöppel et al. 1973; Sanders et al. 1974; Weiskrantz et al. 1974). This phenomenon, which is termed “blindsight,” was first reported in the patients with gunshot wound or vascular lesioning (Pöppel et al. 1973), and then by Weiskrantz and colleagues for a patient with neurosurgical removal of the V1 (Sanders et al. 1974). Since then, a number of articles have been published on the patients with damage to the V1 showing the dissociation between visual awareness and behavioral responses. In most of the human studies, behavioral responses were examined by manual responses such as goal-directed reaching or button press and partly with saccadic eye movements (Danckert and Rossetti 2005; Weiskrantz 2009; Cowey 2010). Furthermore, to clarify the neural basis of blindsight, visuomotor function after V1 lesioning has been tested in non-human primates, either using a manual response task (Humphrey 1974; Feinberg et al. 1978; Cowey and Stoerig 1995) or saccades (Mohler and Wurtz 1977; Segraves 1987; Moore et al. 1995; Schmid et al. 2010) as effectors. More recently, our laboratory has conducted a series of studies to clarify the visual pathways underlying blindsight (Yoshida et al. 2008; Kato et al. 2011, 2021; Kinoshita et al. 2019; Takakuwa et al. 2021) and the visuomotor/cognitive abilities retained in the blindsight condition in the macaques (Yoshida et al. 2008; Ikeda et al. 2011; Yoshida and Isa 2015; Yoshida et al. 2017). In these studies, we mainly used visually guided saccades as the effector, because the circuits for the saccadic motor system were relatively well understood (Krauzlis 2005) and also because a number of behavioral tasks were established with eye movements in macaques. These studies have shown that monkeys with a V1 lesion can perform not only a simple reflexive reaction-time saccade task but also more complex tasks that require cognitive processes such as short-term memory (Takaura et al. 2011) or associative learning including Pavlovian conditioning (Takakuwa et al. 2017) and instrumental learning (Kato et al. 2021) (for review, see Isa and Yoshida 2021). However, the level of conscious control varies depending on the effector. One can imagine that saccades are more reflexive and would not require a high level of conscious perception compared to goal-directed forearm reaching or manual button press tasks. Thus, the experimental settings in previous studies that adopted saccades may not always be the most appropriate for testing the cognitive functions retained in the blindsight condition. In particular, we demonstrated previously that the superior colliculus (SC) is a critical node for blindsight; however, it is also known to be the center of oculomotor control (Gandhi and Katnani 2011; Basso et al. 2021; Isa et al. 2021). Therefore, there is a concern that previous findings that the SC is critical for blindsight might not fully explain the results of previous human patient studies that employed manual response tasks (Perenin et al. 1978; Kentridge et al. 1999). In the present study, as the first step to address these issues, we assessed the ability of V1-lesioned macaques to perform a conditioned manual response task and examined their visuomotor processing by recording the activity close to the motor output stage, tracing back from the primary motor cortex (M1). For this purpose, we recorded the activity of ipsilesional frontal cortical areas using chronically implanted multichannel electrocorticography (ECoG) electrodes to investigate how these areas are involved in visuomotor processing under the blindsight condition. First, we focused on detecting the visual responses related to successful task performance and then analyzed whether these visual responses were transferred to the M1 to generate motor commands using Granger Causality analysis. We found that the dorsal premotor cortex (PMd) plays a critical role in visuomotor transformation in the blindsight condition.

## Methods

### Animals

Two adult macaque monkeys (Macaca fuscata; both male; body weight, 7.5– 8.0 kg [monkey U]; 3.8–4.0 kg [monkey S]) were used in this study. All experimental procedures were performed in accordance with the National Institutes of Health Guidelines for the Care and Use of Laboratory Animals and were approved by the Committees for Animal Experiments at the Graduate School of Medicine in Kyoto University and at the National Institute of Natural Sciences.

### Behavioral task

Psychophysics Toolbox version 3 (http://psychtoolbox.org/) on MATLAB (MathWorks) was used for stimulus presentation and data acquisition. A monitor (Diamondcrysta WIDE RDT272WX [BK]; Mitsubishi] was placed at 600 mm from the eyes of the monkeys. Eye movements were recorded by an eye tracker (EyeLink 1000 PLUS; SR Research) at a sampling rate of 1,000 Hz. All statistical analyses were performed using MATLAB software (MathWorks).

To assess visuomotor function, the monkeys were trained to perform a 2-alternative forced choice manual response (2AFCMR) task (Fig. 1). The subjects were seated in a monkey chair in front of the monitor screen with their heads fixed using a head post, and were required to put their right hand on a lever positioned at the height of their abdomen. In our 2AFCMR task, an initial FP (a donut shape with 1.2° outer and 0.6° inner diameters) was presented at the center of the screen. The monkeys had to move their eyes to the fixation window (3.9° diameter) within 2 s after its presentation and maintain their gaze in the window for 0.5–1.0 s. Another visual stimulus (cue, 0.8° diameter) was then presented randomly at one of two possible locations, each radially separated by 30° relative to the FP (centered [0°] on the horizontal meridian; +15° up or −15° down], in either the left or right visual hemifield (Target cue). After a 0.5–1.0 s delay from cue presentation, the color of the FP changed from red to green, which instructed the monkeys to “go” for the action (Go cue). The monkeys were required to push the lever if the cue was on the upper side of the screen, and pull the lever if the cue was on the lower side of the screen. The luminance Michelson contrast of the FP was 0.95 on a background of 1.0 cd/m^2^. The luminance contrast of the Target cue was systematically changed for monkey U before V1 lesioning; however, it was fixed at a Michelson contrast of 0.95 after V1 lesioning in monkey U and before and after V1 lesioning in monkey S. Water was delivered as a reward if the monkeys made the correct action within a response time of 0.5 s (“Hit” trial). If the monkeys just maintained fixation and made no manual action within the response time, it was judged as a “Miss” trial. In addition, if the monkeys moved the lever in the wrong direction, it was judged as an “Error” trial. Hit, Miss, and Error trials were all included as “valid trials.” If the monkeys did not fixate on the FP, did not maintain fixation until the Go cue, or moved the lever before the Go cue, the trial was regarded as an “invalid trial” and removed from the analysis. If the preceding trial was a Hit trial, the inter-trial interval was set at 0.5 s, but otherwise the interval was set at 2 s. The Hit rate was calculated by dividing the number of Hit trials by the number of valid trials during the experimental block, which consisted of 220 trials with approximately 2 min intervals between the blocks. Two to six blocks were conducted on each experimental day. We only analyzed the trial blocks in which the ratio of valid trials was higher than 70%.

**Figure 1.**
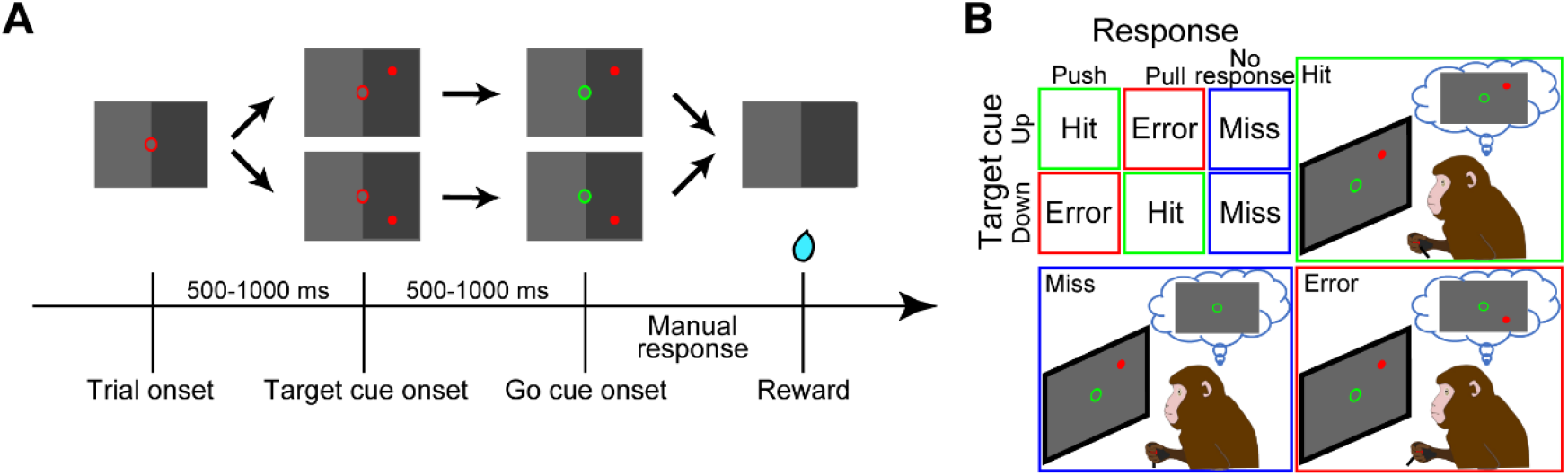
Two-alternative forced choice manual response (2AFCMR) task. (A) The sequence of the stimuli presented in a trial of the 2AFCMR task. (B) Relationship between Target cue location and the required manual response.

### Surgery

A head post was attached to the monkeys under anesthesia induced with xylazine hydrochloride (2 mg/kg) and ketamine hydrochloride (5 mg/kg) and maintained with isoflurane (1.0–1.5%) from a ventilator and intramuscular injection of ketoprofen (0.2 mL/kg). Postoperatively, we treated the animals with intramuscular analgesics (5 mg/kg ketoprofen) and antibiotics (5 mg/kg ampicillin) for 5 days. After 2–6 months of training on the 2AFCMR task, ECoG implantation surgery was undertaken with the same anesthetic protocols as above. Craniotomies were made around the central sulcus, and the cerebral cortical surface around the central sulcus was exposed on the left side. Two platinum ECoG arrays, comprised of multiple channels (48 channels, 6 × 7 + 3 × 2 grids for monkey U [Figs. 4A, 5A, 6A and 7A] and 36 channels, 5 × 6 + 3 × 2 grids (Unique Medical Co., Ltd., Tokyo) for monkey S [Fig. 4L, 5L and 7C]), were placed on the frontal eye field (FEF), premotor cortex (PM), M1, and primary sensory cortex (S1). Each electrode had a diameter of 2 mm and the distance between the center of two adjacent electrodes was 3 mm. The impedance of all electrodes was <100 kΩ. The position of the ECoG arrays was confirmed by combining magnetic resonance images taken before surgery and computed tomography images taken after surgery, and being exposed at the end of all the experiments after transcardial perfusion with 4% paraformaldehyde and photographed. The electrode locations were thus reconstructed (Figs. 4A, 4L, 5A, 5L, 6A, 7A and 7C).

**Figure 2.**
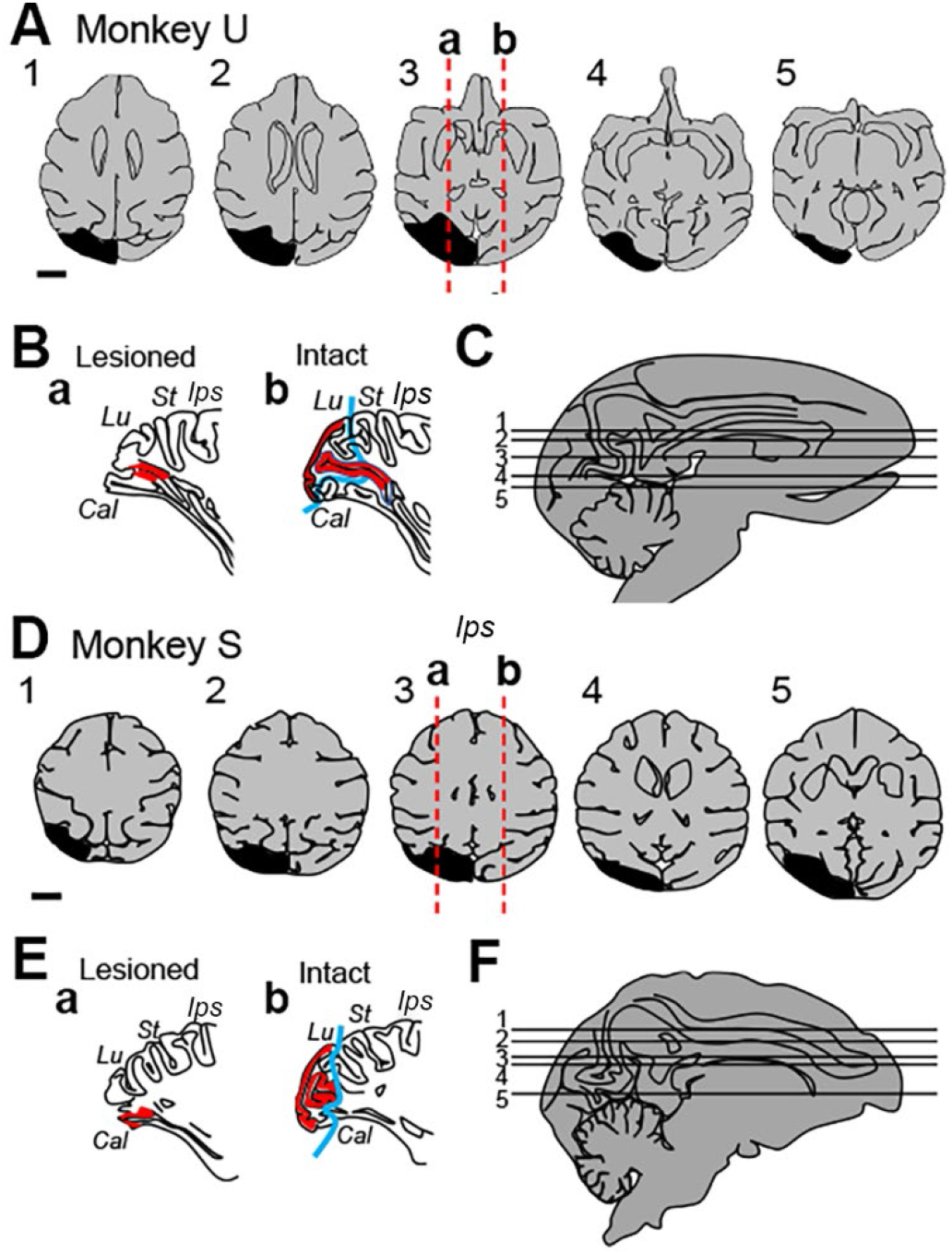
The extent of the primary visual cortex (V1) lesions of monkeys U (**A-C**) and S (**D-F**). In **A** and **C**, the lesioned area is depicted as black shade in the horizontal views. The sagittal views in **Ba** and **Ea** show the lesioned area in the occipital cortex, whose laterality is indicated in **A3** and **D3** with the vertical red dotted line (**a**), respectively. The sagittal views in **Bb** and **Eb** show their counterpart on the intact side, whose laterality is indicated in **A3** and **D3** with the vertical red dotted line (**b**), respectively. Blue lines in **Bb** and **Eb** indicate the rostral border of the lesioned area reflected on the intact side. The extent of the V1 is indicated in red. The individual horizontal sections (1–5) in **A** and **D** are matched in **C** and **F** with corresponding numbers. Abbreviations. Cal: calcarine sulcus, Ips: intraparietal sulcus, Lu: lunate sulcus, St: superior temporal sulcus.

**Figure 3.**
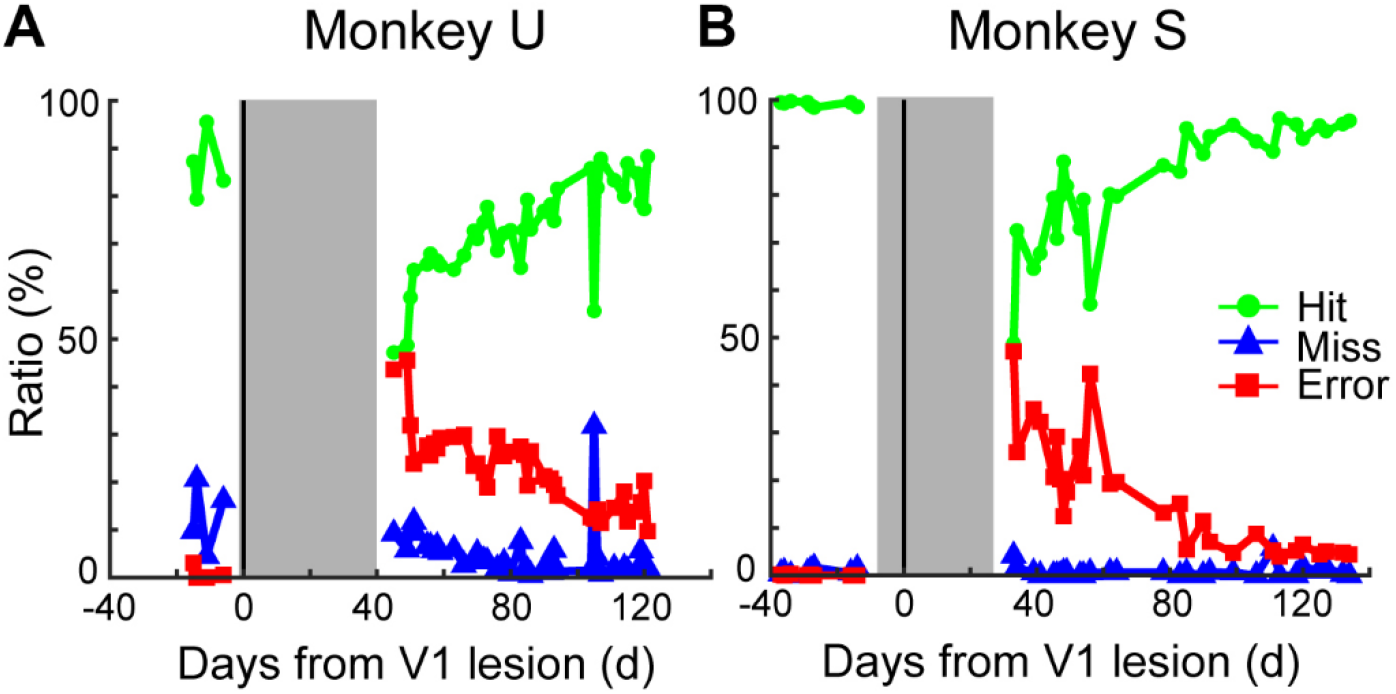
Behavioral performance of monkeys U (A) and S (B). Green, blue and red line indicate the Hit, Miss and Error rates, respectively. Gray area after lesioning indicates the time period in which monkey could not perform the task due to lesioning.

**Figure 4.**
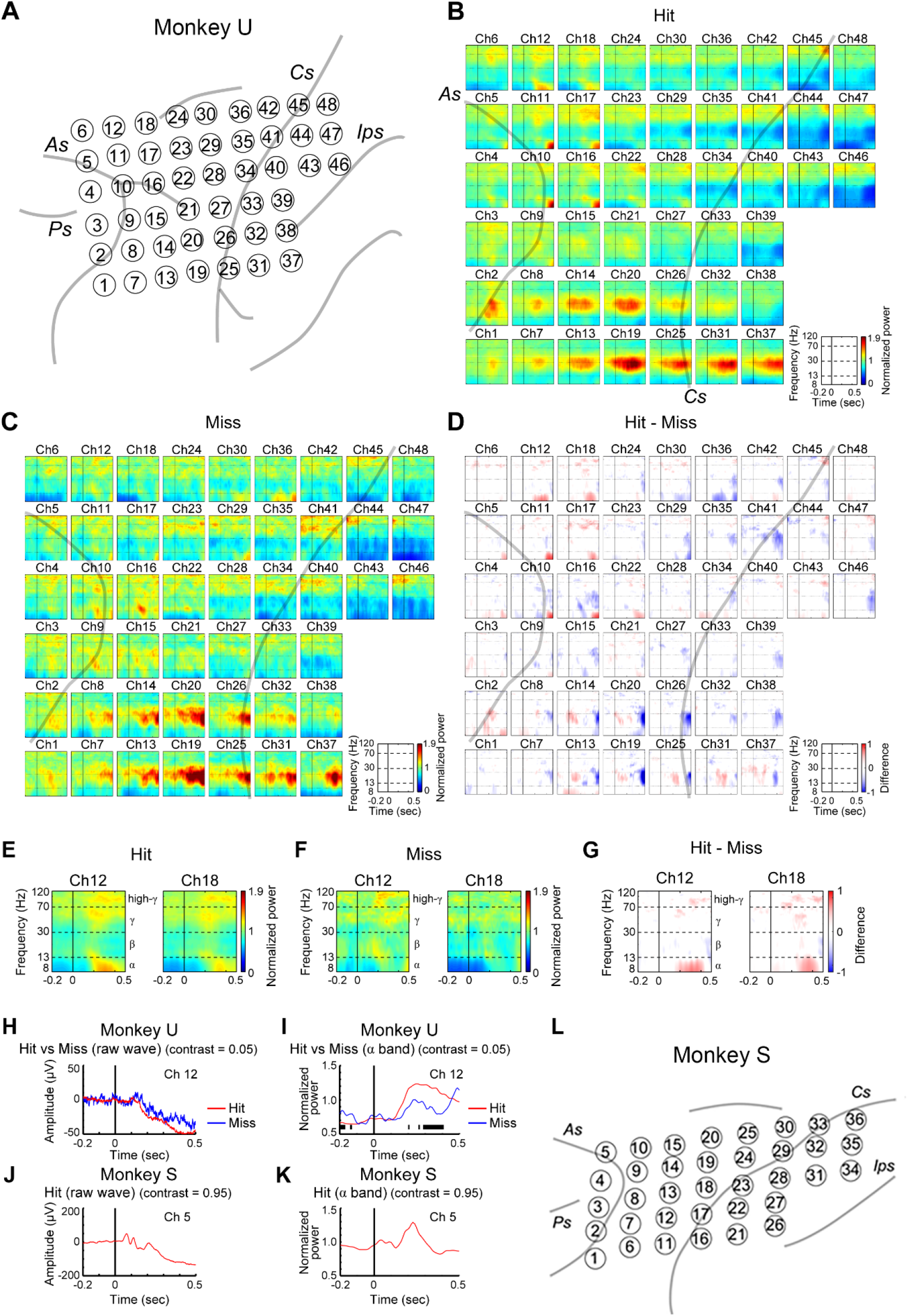
Visual responses before V1 lesioning. (A) Schematic diagram of electrocorticography electrode alignment in monkey U. Abbreviations. As: arcuate sulcus, Cs: central sulcus, Ips: intraparietal sulcus, Ps: principal sulcus. Averaged time-frequency spectra of the Hit (B), Miss (C), and Hit-Miss (D) trials are shown for each channel. The vertical lines corresponding to time 0 ms are aligned on the timing of Target cue presentation. Among these, records from Ch12 and 18 in the dorsal premotor cortex are picked up for the Hit (E), Miss (F), and Hit-Miss (G), respectively. Averaged raw waveform (H) and normalized θ ~ α band activity (I) of Hit (red) and Error (blue) trials around cue onset for Ch 12 in monkey U. Black vertical bars indicate the time points at which a statistically significant difference (p < 0.05) was detected between the two recordings. Averaged raw waveform (J) and normalized θ ~ α band activity (K) around cue onset of the Hit trials for Ch 5 in monkey S. (L) Schematic diagram of electrocorticography electrode alignment in monkey S. Abbreviations. AS: arcuate sulcus, CS: central sulcus, Ips: intraparietal sulcus, PS: principal sulcus.

**Figure 5.**
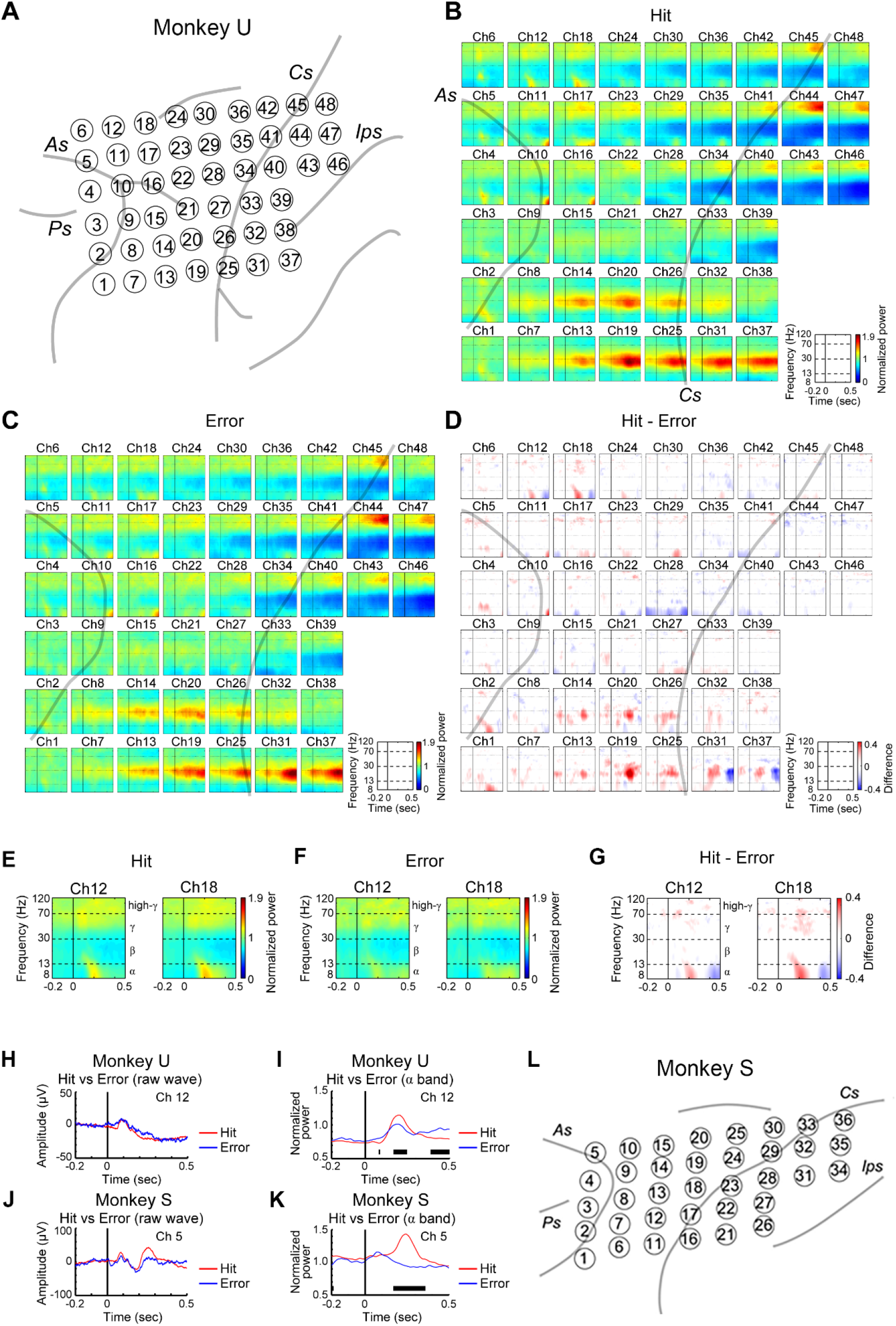
Visual responses after V1 lesioning. (A) Schematic diagram of electrocorticography electrode alignment in monkey U. Averaged time-frequency spectra of the Hit (B), Error (C), and Hit-Error (D) trials are shown for each channel. The vertical lines corresponding to time 0 ms are aligned on the timing of Target cue presentation. Among these, records from Ch12 and 18 in the dorsal premotor cortex are picked up for the Hit (E), Error (F), and Hit-Error (G), respectively. Averaged raw waveform (H) and normalized θ ~ α band activity (I) of Hit (red) and Error (blue) trials around cue onset for Ch 12 in monkey U. Black vertical bars indicate the time points at which a statistically significant difference (p < 0.05) was detected between the two recordings. Averaged raw waveform (J) and normalized θ ~ α band activity (K) around cue onset of the Hit (red) and Error (blue) for Ch 5 in monkey S. (L) Schematic diagram of electrocorticography electrode alignment in monkey S.

**Figure 6.**
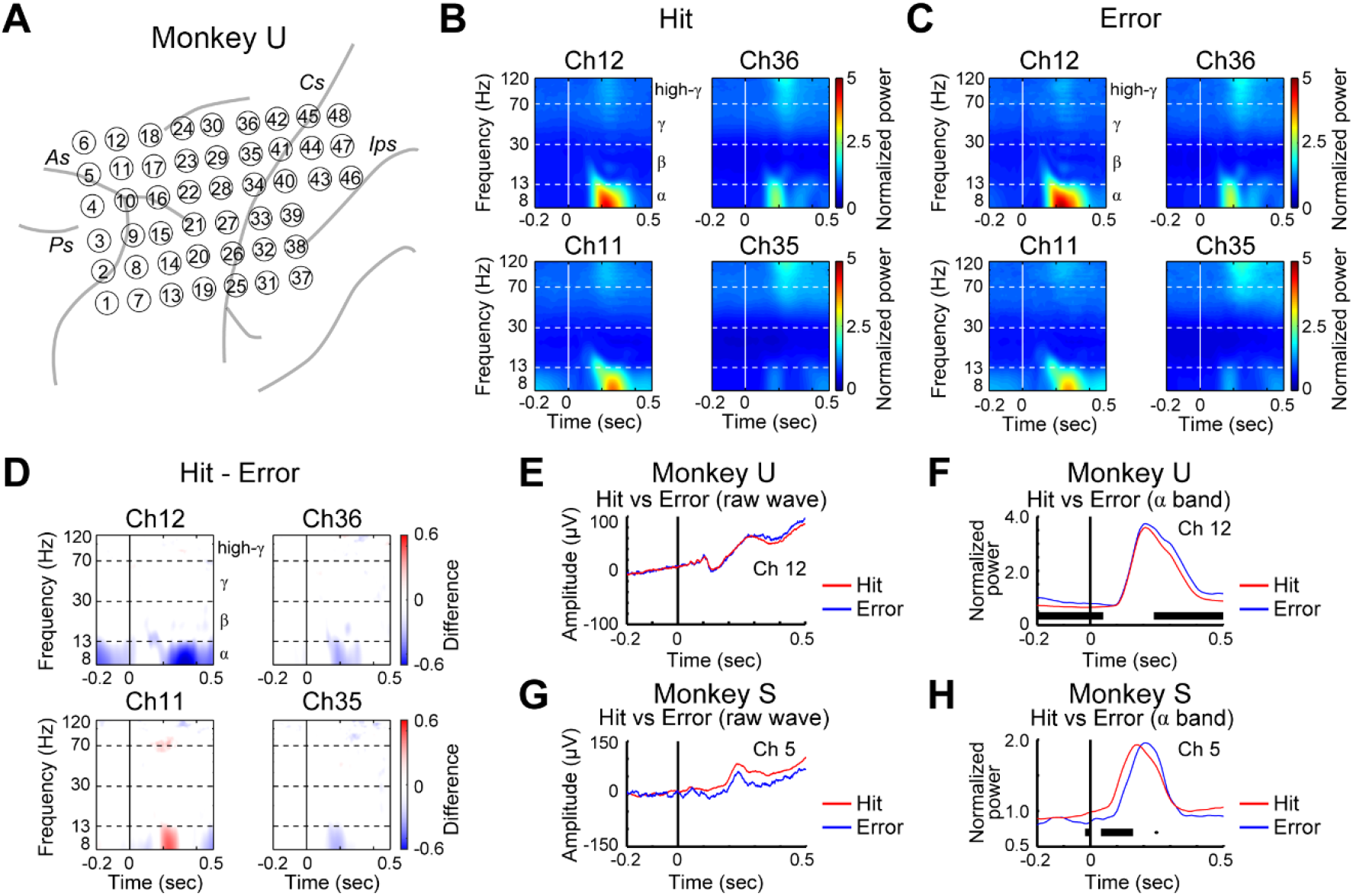
Task-related activity in channels (Ch) 11, 12, 35, and 36 of the electrocorticography (ECoG) electrodes after primary visual cortex lesioning in monkey U. (A) Schematic diagram of ECoG electrode alignment. Averaged time-frequency spectra for the Hit (B), Error (C), and Hit-Error (D) trials are shown for each channel. The vertical lines corresponding to time 0 ms are aligned on the timing of Go cue presentation. Raw wave (E and G) and normalized power of the α band (F and H) for the Hit (red) and Error (dark blue) trials in monkeys U (E and F) and S (G and H). Black bars indicate the time points at which a statistically significant difference (p < 0.05) was detected between the two recordings.

**Figure 7.**
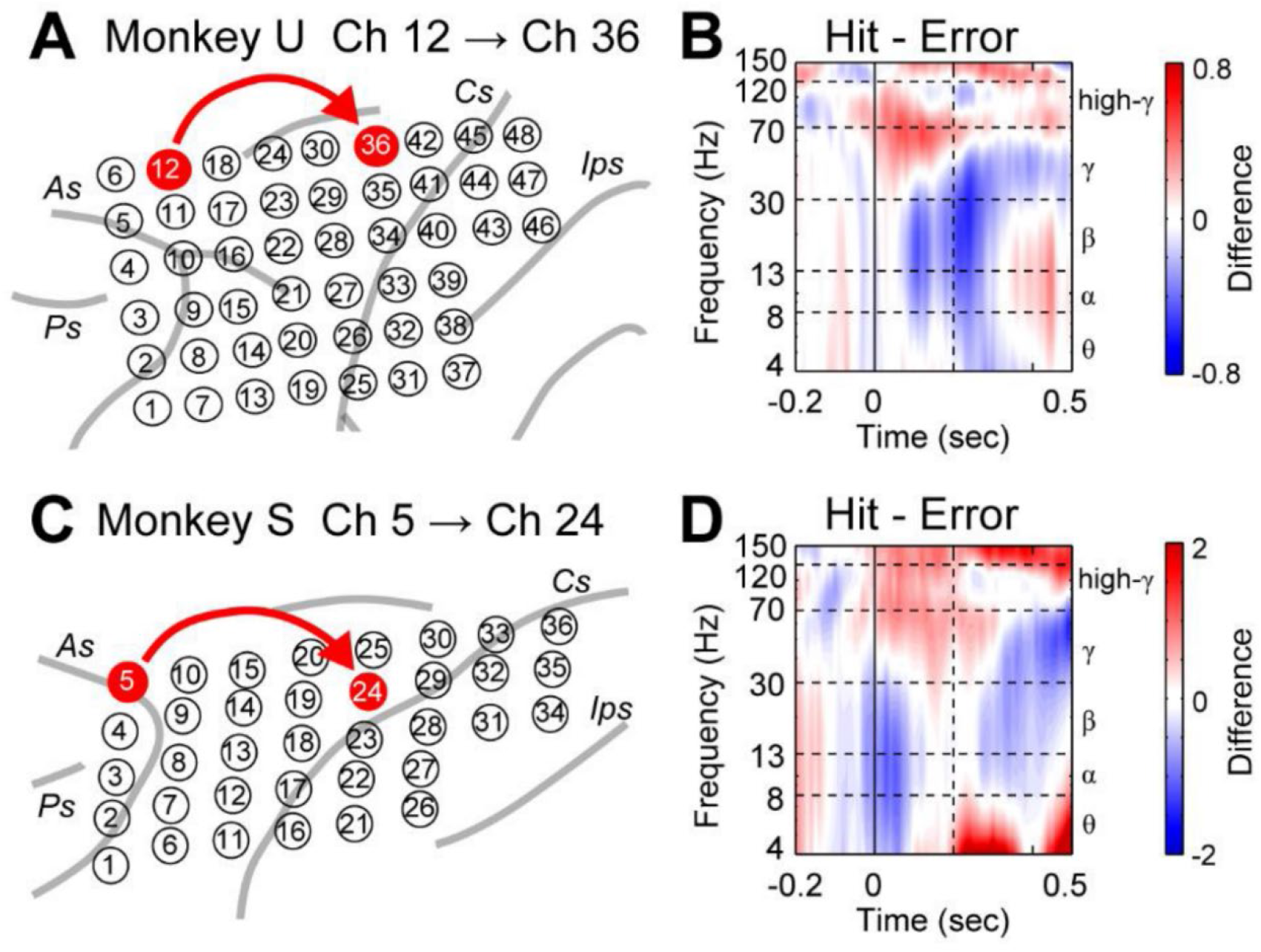
Granger causality analysis from the dorsal premotor cortex (PMd) to primary motor cortex (M1) in the blindsight condition in monkeys U and S. (A) Schematic diagram of electrocorticography electrode alignment in monkey U. Channels (Ch) 12 and 36 were selected as representative channels for the PMd and M1, respectively. (B) Subtraction of Granger causality in the time-frequency dimension around cue onset between the Hit and Miss trials. (C and D) The same arrangement as (A and B), but for monkey S. Ch5 and 24 were selected as representative channels for the PMd and M1, respectively.

After recording the pre-lesioning data, surgery was performed to lesion the V1 unilaterally with the same anesthetic protocols as above. Details of the surgery have been described in previous articles (Yoshida et al. 2008; Takakuwa et al. 2022). In brief, the left V1 of each monkey was surgically removed by aspiration under the anesthesia described above. As shown in Fig. 2, the caudal surface of the V1, except its ventrolateral part, which represents the foveal region, was removed. Furthermore, the caudal aspects of the calcarine sulcus were also aspirated. The lesion also partly included the neighboring V2 and underlying white matter. The behavioral data in the present study were obtained for 4 months (monkey U) and 5 months (monkey S) after V1 lesioning.

### Electrophysiological recording and analysis

ECoG signals were recorded with a data acquisition system (Cerebus; Blackrock Microsystems) at a sampling rate of 1,000 Hz. ECoG signals were extracted using multichannel amplifiers with an analog filter (0.3-Hz high-pass and 7,500-Hz low-pass). For preprocessing, 60-Hz line noise was removed from the raw ECoG data using the MATLAB FieldTrip Toolbox (Oostenveld et al. 2011). The data were then downsampled to 500 Hz. The trials with abnormal spectra were rejected using an automated algorithm from the EEGLAB library (Oostenveld 2011), which has been suggested as one of the most effective methods for artifact rejection. The ECoG signals from each channel were then aligned using the timing of each task event onset (see Figs. 4 and 5). The dynamic properties of cortical activation in individual channels were quantified by time-frequency analysis. We applied multitaper - based time-frequency transformation for the frequency range 1 - 120 Hz by using the open source MATLAB toolbox FieldTrip. Each time-frequency analysis value represented the in-trial dynamics from a channel between −1.0 s and 1.0 s between 1 Hz and 120 Hz. To quantify event-related activity during the task, we further divided each time-frequency analysis value by the baseline value (the mean time-frequency analysis value at the corresponding frequency during the inter-trial interval). To provide a stable representation of in-trial dynamics, we averaged event-related activity across trials collected every 2 weeks. The trial average was calculated separately for the data from before and after lesioning.

After V1 lesioning, ECoG data recordings were continued until the signal to noise ratio deteriorated, which made the analysis difficult in monkey U (18 weeks post-lesioning). In this study, data during the period where performance seemed to have reached a plateau after lesioning were used for analysis. The onset latency of the Target cue response, at which the afferent input arrived in the recording site, was estimated as the time of peak amplitude in the average raw wave in the ECoG recording after the Target cue. The onset latency of the task-related activity was estimated as time in which the difference between Hit and Miss/Error was significant in each bin (Wilcoxon rank sum test, p < 0.05) for the first time after the time 0 ms.

### Granger causality analysis

To examine signal flow from the PMd to M1, Granger causality analysis was conducted. The ECoG data were preprocessed by the MATLAB FieldTrip Toolbox. First, the 60 Hz line noise was removed from the raw ECoG data and the signal was downsampled from 1,000 to 500 Hz. Then, we conducted detrending, temporal normalization, and ensemble normalization of the temporal data. Event-related causality was further quantified by dividing each Granger causality value by the baseline value (the mean Granger causality value at the corresponding frequency during the resting period from −200 ms to 0 s). The model order, which is related to the length of the signal in the past that is relevant to the current observation, was determined by the Akaike information criterion (Akaike 1974). In both subjects, a model order of 10 samples (equivalent to 10 × 4 = 40 ms of history) resulted in a minimum Akaike information criterion and was selected. To illustrate the temporal profile of Granger causality, we used a sliding window with a total width of 120 ms and step width of 10 ms.

### Identification of lesion extent

The extent of the V1 lesion was first assessed by T2-weighted imaging with a 3T magnetic resonance imaging scanner (Verio; Siemens) at 2 weeks after surgery. After the experiments were terminated, monkey U was anesthetized deeply with an intravenous injection of sodium pentobarbital (50–100 mg/kg) and perfused transcardially with 0.05 M phosphate-buffered saline and then with 4% paraformaldehyde in 0.1 M phosphate buffer, pH 7.4. The brain was sectioned for the occipital lobes and placed in sample containers filled with Fluorinert. It was analyzed with a 7T magnetic resonance imaging scanner (Magnetom 7T; Siemens) with knee and insert coils using a 3D gradient echo sequence (TR/TE = 35/10 ms, FA 20° in isotropic 150 μm) for reconstruction of the extent of the lesion. Monkey S was still alive at the time of submission of this article.

## Results

### Behavioral performance

Two macaque monkeys (monkeys U and S) performed a delayed conditioned 2AFCMR task for a liquid reward (Fig. 1A and B). In this task, the monkeys were required to push a lever after presentation of the Go cue (color change of the fixation point [FP]) if the preceding Target cue was presented in the upper visual field relative to the FP, and to pull the lever in response to the Target cue being presented in the lower visual field. In monkey U, before V1 lesioning, the luminance contrast of the Target cue was systematically changed and the test was conducted mainly at the near-threshold intensity (Michelson contrast = 0.05, 4 sessions). In this case, the Hit rate was 86.5% (621/718), Miss rate was 12.5% (90/718), and Error rate was 0.9% (7/718) (Fig. 3A). Conversely, monkey S had difficulty in performing many trials with this task at the same luminance contrast level; therefore, the visual stimuli were presented at the maximum contrast (0.95, 19 sessions). The Hit rate in these sessions was 99.2% (3,595/3,623), Miss rate was 0.7% (24/3,623), and Error rate was 0.1% (4/3,623) (Fig. 3B).

After recording the pre-lesioning data, the left V1 was aspirated in both monkeys (Fig. 2). After lesioning, task performance was tested at the full luminance contrast (0.95) in both monkeys. Initially, the monkeys could not perform the task at all; however, through daily trainings, they became able to perform the task with a >70% “valid trial” rate (in which the animals did not interrupt the task before the Go cue; see Methods) in one session (220 trials) on Day 46 after V1 lesioning in monkey U and Day 34 in monkey S (Fig. 3). After that, task performance recovered gradually and reached a >80% success rate on Day 95 in monkey U and >90% on Day 85 in monkey S, both of which were close to the Hit rate before lesioning. It is difficult to make a direct comparison, but, in previous studies on saccades (Yoshida et al. 2008), the monkeys showed >90% recovery within 2 months. Thus, the recovery of the current manual response task appeared to be slower than the recovery for saccades, presumably reflecting the difficulty of the task.

Notably, changes in behavioral bias were observed in both monkeys. Before lesioning, when monkey U failed to get the reward, it mostly missed the Target cue (Miss), that is, it did not respond to the stimuli presumably because this animal was tested with near-threshold stimuli. In contrast, monkey S did not show Miss trials, because it was tested only at the maximum intensity of Target cue luminance. However, after lesioning, both monkeys tended to make errors (Error), in which they made incorrect reactions to the Target cue presented in the affected field. This suggested that the ability to detect and respond to the Target cue in the affected field was preserved; however, the monkeys could not correctly localize the cue in the error trials.

Before lesioning, the reaction time of the manual response for Hit trials was 306 ± 47 ms (n = 516 trials) in monkey U and 227 ± 50 ms (n = 869 trials) in monkey S. After lesioning and the re-training period when task performance exceeded 80%, the reaction time in the successful trials was 318 ± 32 ms (n = 1,780 trials) in monkey U and 227 ± 37 ms (n = 1,400 trials) in monkey S. Thus, the reaction times before and after lesioning were in a similar range; however statistical analysis revealed that the reaction time after V1 lesioning was significantly longer in monkey U (two-sample t-test, p < 0.05), but not in monkey S (two-sample t-test, p = 0.78).

### Visual responses in the frontal cortex with an intact V1

To investigate the neural basis of visuomotor transformation in this task, we analyzed the visual responses to the Target cue stimuli recorded from the ECoG array (48 channels in monkey U and 36 channels in monkey S) placed on the frontal cortical areas to the side which we planned to make the V1 lesioning (left hemisphere), including the FEF, PM, M1, and S1. We chose the ECoG for recording the cortical activity because it can cover a wide area of the frontal cortical areas and promises the direct comparison between the pre- and postlesion states. Fig. 4H and J show the raw waves of ECoG recording in the Hit (red) and Miss (blue) trials from channel (Ch) 12 and Ch 5 in monkey U and S, respectively. The event-related change in the ECoG response started at 50–80 ms after Target cue onset in the positive direction, while the negative visual response started with a latency of 109 ms for monkey U (contrast 0.05). A similar fast negative visual response was observed with a latency of 70 ms in monkey S (contrast 0.95) (Fig. 4H and J). To analyze the ECoG responses further, time-frequency analysis was conducted. Here, neural activity was normalized through dividing the data from the period of interest by the baseline activity (taken from −200 to 0 ms before Target cue onset). Figs. 4B and C show time-frequency analysis of the neural activity recorded with the ECoG electrodes on the left FEF/PM/M1/S1 of monkey U during performance of the 2AFCMR task with its right arm in the prelesion state. In this monkey, task-related activity (Fig. 4D) was calculated by subtracting the average activity in the Miss trials (Fig. 4C, n = 87 trials) from that in the Hit trials (Fig. 4B, n = 521 trials). The most prominent activity related to task performance was observed in channels in the PMd (Ch12 and 18 in Fig. 4E-G), which showed larger response in the Hit trials than in the Miss trials primarily in the θ ~ α band (4–13 Hz), at 200–500 ms after Target cue presentation (hereafter we term “task-related θ ~ α band Target cue response”, which could not be recorded in the adjacent electrode channels such as Ch11 and 17 (see Fig. 4D). We observed prominent responses in the β band (14–30 Hz) from the more lateral electrodes (Ch13, 14, 19, 20, 25, 31, and 37) in the Hit trials; however, they were mostly eliminated by subtracting the activity in the Miss trials, despite some negative components that remained later than 400 ms after Target cue presentation. Thus, these responses cannot be regarded as meaningful task-related visual responses. Therefore, we focused on the θ ~ α band short latency visual responses in the PMd. Fig. 4I shows the θ ~ α band component derived from the raw record in Fig. 4H. The θ ~ α band component was larger in the Hit trials than in the Miss trials (Wilcoxon rank sum test, p < 0.05) at 288–408 ms after Target cue onset. Similarly, the θ ~ α band component was increased in the PMd of monkey S (contrast 0.95) (Fig. 4K). Monkey S made almost no Miss trials, because strong visual stimuli were used for the Target cue. Therefore, only Hit trials were analyzed before V1 lesioning.

### Visual responses in the frontal cortex after V1 lesioning

After V1 lesioning, the same analysis was conducted as before lesioning (Fig. 5). Here, because of the behavioral change mentioned above, the Error trials were used for comparison with the Hit trials. In the raw wave, the latency of the negative visual response was detected by the initial positive peak at 85 ms for both monkeys (Fig. 5H and J). The difference between the Hit and Error trials was not very obvious; however, time-frequency analysis showed a clear θ ~ α band task-related Target cue response that demonstrated a distinct difference between the Hit and Error trials at approximately 160 ms after Target cue presentation (Fig. 5I and K). In detail, when the monkeys successfully detected and responded to the Target cue, the time-frequency analysis map showed a significant difference in the power of θ ~ α band activity at 168–248 ms after Target cue presentation between the Hit and Error trials in the PMd of monkey U (Ch12 and 18) (Fig. 5B-G and I), and at 168–360 ms in monkey S (Ch5, Fig. 5K) (Wilcoxon rank sum test, p < 0.05), similarly to the pre-lesioning data (Fig. 4). Thus, the task-related θ ~ α band Target cue response was detected at approximately 200 ms after cue presentation in the PMd after V1 lesioning.

### Granger causality from PMd to M1

To investigate whether the task-related visual responses observed in the PMd led to the motor command in the M1, we conducted Granger causality analysis. ECoG recordings after Go cue onset showed a clear increase in γ and high-γ band activity in the electrodes in the M1, which is known to be related to the movement execution (Ch 35 and 36 in Fig. 6. Here, there was no significant difference in the activity after the Go cue onset between the Hit and Error trials as expected). In this analysis, we selected Ch12 and 5 as source channels in the PMd for monkeys U and S, respectively, and Ch36 and 24 as sink channels in the M1 hand area for monkeys U and S, respectively (monkey U, Fig. 7A; monkey S, Fig. 7C). We calculated Granger causality in the Hit and Error trials in the time-frequency dimension and subtracted the latter from the former to visualize the dynamic changes in the patterns of signal flow from the PMd to M1 (Fig. 7B and D). A causal effect from the PMd to M1 was observed in the θ ~ α band (4–13 Hz) during the delay period at 300–500 ms after Target cue presentation in both monkeys (Fig. 7B and D). In addition, Granger causality in the γ band (31–70 Hz) was enhanced during the first 200 ms after Target cue presentation and then declined. The high-γ band (>70 Hz) was enhanced soon after Target cue onset, which continued toward Go cue onset. In contrast, a difference in Granger causality in the β band appeared to be suppressed during the delay period. The results were strikingly consistent across both monkeys. These findings suggest that the PMd contributed to the subsequent activity in the M1 to initiate manual responses to the visual cue in the blindsight condition through multiple frequency channels, especially in the θ ~ α, γ, and high-γ bands.

## Discussion

In this study, we have shown that blindsight monkeys could retain the ability to perform the conditioned manual response tasks in response to Target cue stimuli with arbitrary assignment of the cue-response relationship (upper cue = lever press vs. lower cue = lever pull) presented in the affected visual field through post-injury training over a period of 2– 3 months. At this stage, their success rate reached >80% and manual response reaction times were in a similar range to those in the intact state. One can imagine that it would be difficult to conduct these tasks with a complete loss of awareness of the visual cue. This observation provides further support for our hypothesis that the monkeys had some form of conscious sensing of the visual target, similar to “type II” blindsight patients who describe “feeling something happening” in their affected visual field (Yoshida and Isa 2015; Isa and Yoshida 2021). Furthermore, ECoG recordings revealed that the M1 hand area showed high-γ band activity during the manual response, while the PMd showed θ ~ α band task-related Target cue responses with latencies at approximately 200 ms, both similar to the results observed in the intact state. Granger causality analysis revealed that there was clear signal flow from the PMd to M1 during the delay period in the θ ~ α, γ, and high-γ bands, which would have led to the successful performance of the manual response task. These findings provide new insights in the field of blindsight research.

### Contribution of PMd to visuomotor transformation

A number of previous studies on blindsight subjects have focused on extrastriate visual areas such as the V2, V4 (Cowey and Stoerig 1991; Ajina et al. 2015; Bridge et al. 2019), MT (Schmid et al. 2010; Rodman et al. 1989; Ajina et al. 2018) and LIP (Schmid et al. 2010; Kato et al. 2021) in the context of visual processing that bypasses the V1, mostly using the no-invasive neuroimaging techniques with low temporal resolution except the electrophysiologcal recording in Kato et al. (2021) and Rodman et al. (1989). Conversely, here, we aimed to clarify the visuomotor processing for hand movement control tracing back from the output stage. In the case of lever push-pull task adopted in this study, it is highly likely that the hand area of the M1 primarily controls movements by sending commands to the spinal cord. In intact animals, it is generally considered that the PM mediates the visual signals to M1 for such visually guided manual responses (Jeannerod et al. 1995; Hoshi et al. 2007; Nakayama et al. 2008). However, it was not obvious whether the PM is still responsible for mediating the visual signal in the absence of V1, and even if this is the case, it is not clear which subregion of PM is responsible. Therefore, to reveal the neural substrate for visuomotor processing for hand movement control in the blindsight condition, multichannel recordings with ECoG were conducted widely covering the FEF, PM, M1, and S1 to clarify the visual inputs to the frontal cortex after V1 lesioning and whether the visual signal is utilized to activate the M1 for hand movement control. As a result, we found that, similar to the intact state, the PMd showed movement-related Target cue responses in the θ ~ α band at ~200 ms after cue onset, which was higher in the Hit trials than in the Error trials even after V1 lesioning. The involvement of the PMd in the conditioned motor task has been reported previously in the intact state (Hoshi and Tanji 2007; Nakayama et al. 2008). Interestingly, even without the V1, the same PM area was used to control the conditioned manual responses. The onset latency of the visual response in the PMd was slightly longer after V1 lesioning (~85 ms) than in the intact state (~70 ms). We will discuss the possible input pathway in the later section.

### Signal flow from the PMd to M1

As the next step, we examined the signal flow from the PMd to M1 by Granger causality analysis to obtain evidence supporting the hypothesis that the task-related visual cue response in the PMd was actually transmitted to the M1, and if so, how was the signal transmitted. Our present results showed that time-frequency analysis of Granger causality during the delay period revealed an increase in signal flow from the PMd to M1 in the θ ~ α, γ, and high-γ band ranges, while it was suppressed in the β band. These findings were robustly confirmed in both animals. As shown in Fig. 6, the M1 demonstrated prominent γ and high-γ band activity around movement onset, as reported previously (Aoki et al. 1999). Thus, it was likely that the task-related visual responses in the θ ~ α band were either directly transferred to the M1 or converted to the γ or high-γ band signal in the PMd and then transferred to the M1. In the case of the visual system, Bastos et al. showed through large-scale ECoG recording in awake monkeys that the bottom-up signaling in the visual areas is coded in the θ (4–7 Hz) and γ (60–80 Hz) bands, while the top-down influence is carried by the β band (14–18 Hz) (Bastos et al. 2015). In addition, late (>200 ms from cue presentation) γ band activity is also suggested to contribute to the coupling between the frontal cortex and visual areas such as the V4 (Gregoriou et al. 2009). Compared with the visual system, few studies have examined top-down or bottom-up signal transmission in the motor system; however, if we consider that the PMd to M1 pathway is bottom-up, the same logic as applied to the visual stream may fit with the hierarchy of the motor system (Chao et al. 2019). These results suggest that the PMd functions as a hub for bottom-up visuomotor processing in the frontal cortex in the blindsight state. Its relationship to conscious/unconscious visual processing needs careful inspection in future studies.

### Visual input pathway to the PMd after V1 lesioning

What is the neural substrate for the induction of the task-related visual responses in the PMd after V1 lesioning? In this study, the earliest onset latency of the visual responses to the Target cue stimuli after V1 lesioning (~85 ms) was in a similar range as in the intact state (~70 ms). They are both in a similar range to the reported findings in frontal cortical areas for comparisons of the visual response latencies in multiple brain areas (Schmolesky et al. 1998). It seems that the visual signal was transformed to the task-related signal which emerged in the same region at the latency around 200 ms. Multiple visual pathways have been proposed for blindsight; previous studies suggested that direct projections from the lateral geniculate nucleus to the extrastriate visual area underlie blindsight in nonhuman primates (Cowey and Stoerig 1991; Schmid et al. 2010; Bridge et al. 2019) and humans (Bridge et al. 2008; Ajina et al. 2015, 2018). Conversely, the involvement of the SC to pulvinar pathway has also been proposed (Diamond and Hall 1969; Cowey and Stoerig 1991; warner et al. 2015), and our group obtained direct evidence of the involvement of this pathway by pathway-selective blockade using a double viral vector intersectional technique (Kinoshita et al. 2019). Furthermore, in the case of saccade control, by using reversible pharmacological inactivation, we found that the lateral geniculate nucleus (LGN) and pulvinar play critical roles after recovery from V1 lesioning, while only the LGN is critical in the intact state, which suggested plastic changes to upregulate the SC-pulvinar pathway after V1 lesioning (Takakuwa et al. 2021). In the present study, we showed that the PMd could mediate the visual inputs to the M1 to perform the conditioned manual response task. Then, how could the PMd become a node for visuomotor processing for blindsight? From the analogy of saccade control, the LGN-extrastriate and/or the SC-pulvinar-extrastriate visual pathways mediate the visual signal to the parietal cortex, especially area 7b, which is considered to interact with the premotor cortices for goal-directed hand movement control (Jeannerod et al. 1995; Rushworth et al. 1997). Another possible route might be the subcortical pathway from the SC to the mediodorsal nucleus of the thalamus and then to the frontal areas (Sommer and Wurtz 1998). Previous studies have suggested that monkeys with V1 lesion exhibit deficit in visual awareness of the target according to their behavior in the Yes-No task condition (Cowey and Storig 1995) and its analysis by signal detection theory (Yoshida and Isa 2015). The amount of overlap between the compensatory pathway after V1 lesioning and the visuomotor pathway in the intact state may affect the flexibility and self-efficacy of task control in the blindsight condition. An obvious next step is to investigate the neural pathway to PMd by reversible inactivation of the possible relay stations such as the LGN, SC, pulvinar, and mediodorsal nucleus of the thalamus and testing its effect on the neural responses in the PMd and behavioral outcomes.

## Author Contributions

YY, SK, JT, HO, RY, and TI designed the experiments and prepared and conducted the surgery necessary for the experiments. YY, SK, and RY conducted the recording experiments and analyzed the data. YY, SK, and TI mainly wrote the manuscript. All authors confirmed the final draft of this article.

## Acknowledgments

We thank Norihiro Takakuwa for comments on earlier version of the manuscript, Zenas C. Chao for instruction on the ECoG data analysis, Jiro Yamashita, Yuta Shinto, Masashi Nakamura, Atsuna Mori and Kei Kubota for technical assistance. We thank Tomohisa Okada and Shin-ichi Urayama for performing ex-vivo MRI of the postmortem brain. This study was supported by grants from CREST of JST (Grant No. JPMJCR1651), KAKENHI from the Ministry of Education, Culture, Sports, Science, and Technology of Japan (26221003, 19H01011, 19H05723) and Brain/MINDS-Beyond of Japan Agency for Medical Research and Development (20dm0307005h0003) to T.I. An earlier version of the draft of this manuscript was edited by NAI, Inc. (Yokohama, Japan).

